# Increased male investment in sperm competition results in reduced maintenance of gametes

**DOI:** 10.1101/2022.03.18.484900

**Authors:** Mareike Koppik, Julian Baur, David Berger

**Affiliations:** Department of Ecology and Genetics, Animal Ecology, Uppsala University, 75236 Uppsala, Sweden; Department of Zoology, Animal Ecology, Martin-Luther University Halle-Wittenberg, 06120 Halle (Saale), Germany

**Keywords:** sexual selection, sperm competition, experimental evolution, mutation rate, post copulatory, DNA repair, oxidative damage, genetic variation, offspring quality, genetic load

## Abstract

Male animals often show higher mutation rates than their female conspecifics. A hypothesis for this male-bias is that competition over fertilization of female gametes leads to increased male investment into reproduction at the expense of maintenance and repair, resulting in a trade-off between male success in sperm competition and offspring quality. Here, we provide evidence for this hypothesis by harnessing the power of experimental evolution to study effects of sexual selection on the male germline in the seed beetle *Callosobruchus maculatus*. We first show that 50 generations of evolution under strong sexual selection, coupled with experimental removal of natural selection resulted in males that are more successful in sperm competition. We then show that these males produce progeny of lower quality if engaging in socio-sexual interactions prior to being challenged to surveil and repair experimentally induced damage in their germline, and that the presence of male competitors alone can be enough to elicit this response. We identify 18 candidate genes that showed differential expression in response to the induced germline damage, with several of these previously implicated in processes associated with DNA repair and cellular maintenance. These genes also showed significant expression changes across socio-sexual treatments of fathers and predicted the reduction in quality of their offspring, with expression of one gene also being strongly correlated to male sperm competition success. Sex differences in expression of the same 18 genes indicate a substantially higher female investment in germline maintenance. While more work is needed to detail the exact molecular underpinnings of our results, our findings provide rare experimental evidence for a trade-off between male success in sperm competition and germline maintenance. This suggests that sex-differences in the relative strengths of sexual and natural selection are causally linked to male mutation bias. The tenet advocated here, that the choices of an individual can affect plasticity of its germline and the resulting genetic quality of subsequent generations, has several interesting implications for mate choice processes.

## Background

The germline mutation rate impacts on a range of evolutionary processes such as the rate of adaption [1, 2] and risk of extinction [3, 4], as well as genome evolution [5]. Contrary to the typical assumption in population genetic models, recent studies have shown that mutation rate can vary both within and between individuals within a given species [6–16]. Such variability can, for example, affect genetic load at mutation-selection balance [17, 18], mate choice and the efficacy of sexual selection [7], and adaptation in stressful environments [8, 19, 20]. Despite these important implications, experimental evidence providing ultimate causation for the observed intraspecific variability in mutation rates remains scarce [21–23].

One prominent type of intraspecific variation is that between males and females of a given species. Males often show higher germline mutation rates in animal taxa [24–28], including humans [29, 30] and other primates [22, 28]. This male mutation bias has been ascribed to the greater number of cell divisions occurring in the male germline prior to fertilization, and the higher number of divisions in males is itself thought to be a result of anisogamy and sexual selection promoting increased gamete production in the sex competing most intensively for fertilization success [31, 32]. Indeed, a need for fast-dividing male germline cells would inevitably lead to an elevated risk of unrepaired replication errors in male gametes, all else equal, as the DNA-repair system must constantly attend single and double strand breaks that occur during meiosis and mitosis [33–36] and in post-meiotic chromatin remodelling during spermiogenesis [29, 36, 37]. This should result in a trade-off between increased male germline replication rates, granting greater success in sperm competition, and increased germline mutation rate, reducing offspring quality [7, 33, 38–41].

However, sex differences in the number of germline cell division do not perfectly predict male mutation bias across species [22, 25, 27, 42, 43] and in humans differences in mutation rate between males of the same age can be many times greater than that between the sexes [12, 13, 30]. Both points suggest that maintenance processes are central in deciding the germline mutation rate. Maintenance of the germline is energetically costly [36, 37, 44] and comprises interrelated processes such as antioxidant defence [45–47], repair of DNA damage [20, 34] and programmed cell death of damaged sperm [48]. Indeed, the male gonad is a highly oxidative environment that, without antioxidant defence devoted to dealing with reactive oxygen species [41, 45–48], produces DNA damage that may result in germline mutations [35–37, 49].Recent studies imply that costly male allocation decisions involving sperm and ejaculate production and composition [50–56] are responsive to female characteristics such as mating status [57] and to the presence of conspecific males [52, 58], most likely because these serve as cues for predicting mating opportunities and the level of competition a male’s sperm may encounter [59]. Thus, socio-sexual cues that signal increased risk of sperm competition and increase allocation to ejaculate components that increase male postcopulatory reproductive success could lead to concomitant plastic decreases in germline maintenance and reduced gamete quality. However, direct experimental evidence supporting this hypothesis remains very scarce indeed.

Here we tested this prediction using experimental evolution lines of the seed beetle *Callosobruchus maculatus*, a model organism for sexual selection where sperm competition is rife [7, 60–64]. These lines have been maintained for >50 generations under three alternative mating regimes manipulating the relative strength of natural and sexual selection: natural polygamy applying both natural and sexual selection (N+S regime), enforced monogamy applying natural selection while removing sexual selection completely (N regime), or a sex-limited middle class neighborhood breeding design [65, 66] applying sexual selection while minimizing natural selection on fecundity and juvenile viability (S regime). The S regime does not show any strong signs of decline in fitness-related traits when assayed at standard conditions [67, 68]. However, previous findings suggest that S-males pass on a greater genetic load to their progeny if having engaged in socio-sexual interactions prior to being challenged with a dose of irradiation introducing DNA-damage in their germline, an evolved response to socio-sexual interactions not seen in N- or N+S-males [7]. A plausible explanation for this result is that S-males have evolved reduced germline maintenance as a response to increased post-copulatory sexual selection coupled with weakened constraints on the evolution of sperm and ejaculate traits in this mating regime, where viability selection was minimized.

In order to test this hypothesis, we first determined sperm competitiveness in males of all experimental evolution regimes to confirm that S males indeed evolved adaptations to the increased post-copulatory sexual selection. Subsequently, we focused on the socio-sexual effect on germline maintenance in S males, as those were the only ones that showed a response to the socio-sexual setting. Here, we set out to determine if male presence, female presence or both triggered the change in germline maintenance in S males. Additionally, we employed RNA sequencing of the reproductive tracts of S males to gain insight into the possible mechanisms behind this change. In a last step, we compared the expression of irradiation responsive genes in males and females (of all experimental evolution lines), as male mutation bias is one of the biggest sources of interspecific variability in mutation rates, and gene expression may shed light on a possible proximate causation.

## Results

### Sperm competition

To test whether the removal of natural selection in S-males has led to the evolution of traits ensuring higher post-copulatory competition success, we assayed male sperm competition success in defense (P1: focal male is first to mate) and offense (P2: focal male is second to mate) for all three evolution regimes, three lines for N and N+S and two for the S regime (one line was accidentally lost during the experimental evolution). Females from the ancestral population, from which the experimental evolution lines were derived, were mated twice (once to a focal male and once to a competitor) with 24 h in between matings, during which time the females were provided with beans for egg laying. The competitor males were from a black mutant strain [69] such that paternity in offspring could be determined. Focal males were either held singly or in groups of five males from the same line prior to mating, and for sperm offense males were also tested in 5 consecutive matings to determine sperm and seminal fluid depletion patterns. Sperm competition success was highest in S males for both, sperm offense and defense (Fig. 1). S males had a significantly higher overall sperm competition success compared to N males (P_MCMC_ = 0.018) and also a strong tendency for higher success compared to N+S males (although this effect was marginally non-significant, P_MCMC_ = 0.078). We found no indication that the advantage of S males depended on being previously exposed to male competitors or having been mated repeatedly prior to the trial, although statistical power was low due to large variation between experimental blocks. Similarly, the sperm competition advantage of S males was not significantly different for offense and defense (Fig. 1, for detailed statistics see Supplementary Information S2).

**Figure 1:**
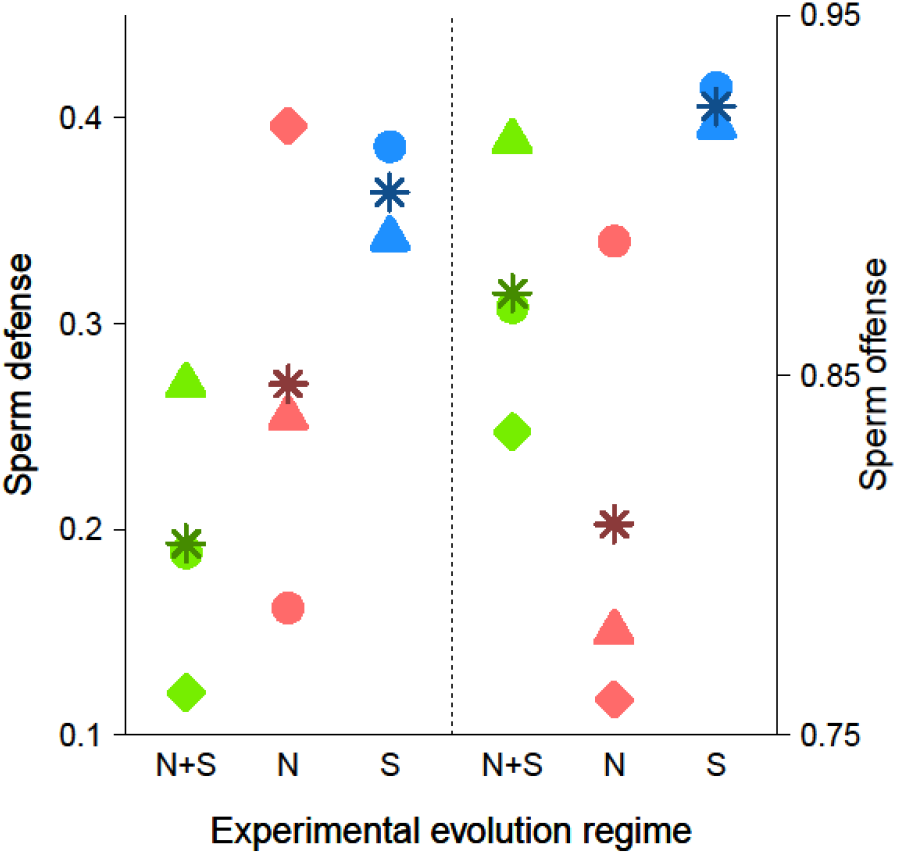
Sperm competition success in males from the experimental evolution lines. Line specific sperm defense (paternity share when male is first to mate, P1) and offense (paternity share when male is second to mate, P2) success in double mated females. Asterisks depict means of experimental evolution regimes. Throughout the manuscript triangles refer to S1, N1, N+S1, diamonds refer to N2, N+S2 and circles refer to S3, N3, N+S3.

### Germline maintenance

#### Offspring quality

Since S males evolved enhanced post-copulatory competitiveness, we hypothesized that they invest more into mating and competition than N and N+S males at a potential cost of reduced germline maintenance. Indeed, S males have previously been shown to reduce germline maintenance when engaging in inter-and intrasexual interactions with conspecifics (Fig. 2A and Baur and Berger (2020) [7]). To dissect the effects of inter- and intrasexual interactions on germline maintenance, we manipulated the social environment of S males in a full-factorial design (with or without male competitors and with or without female mating partners, Fig. 2B). We then measured the reduction in offspring quality for those males after a short (~ 3h) and long (~24h) recovery period after induction of germline damage through gamma radiation. To this end, we mated males to a single virgin female at each time point (3h and 24h post the irradiation treatment) and established a second generation from the resulting offspring. This allowed us to estimate the quality of offspring from F0 irradiated fathers by counting the number of F2 progeny produced in those lineages relative to F2 progeny production in lineages deriving from unirradiated F0 control males. Hence, the reduction in offspring quality could be calculated as: 1-[F2_IRRADIATED_/F2_CONTROL_].

**Figure 2:**
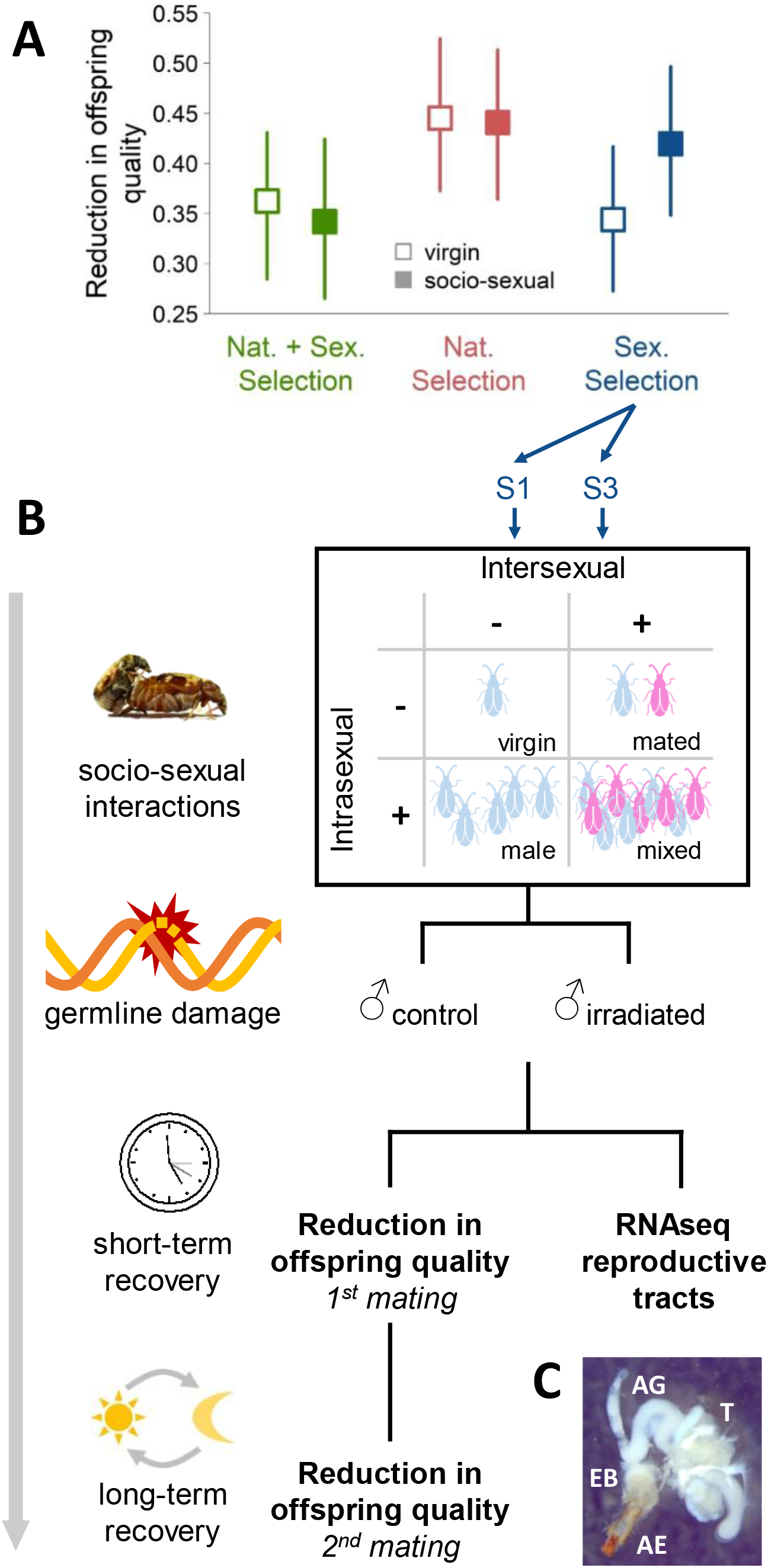
(A) Reduction in offspring quality after induction of germline DNA damage through irradiation of male beetles. Virgin males (open symbols) with an evolutionary history of sexual selection (N+S- and S-males) suffer less reduction in offspring quality than males from lines with only natural selection acting (N-males). However, socio-sexual interactions prior to the challenge to the germline decreases offspring quality further in S-males but not in N+S- and N-males (closed symbols). Data represent Bayesian posterior means and 95% confidence intervals from Baur and Berger (2020) [7]. (B) Schematic overview of the experiment estimating germline maintenance. Males from two S-lines were exposed to one of four socio-sexual environments, manipulating the presence of conspecific males and females. Afterwards, we induced germline damage via gamma radiation and determined reduction in quality of offspring produced by those males after a short (~3h) and long (~24h) recovery period. Additionally, we examined gene expression in male reproductive tracts at the end of the short recovery period. (C) Picture of a male reproductive tract. In *C. maculatus* the male reproductive tract consists of the aedagus (AE), ejaculatory bulb (EB), five accessory gland (AG) pairs (two large and three small AG pairs) and a pair of bilobed testes (T). For the gene expression the two large AG pairs were not included.

In agreement with previous results [7], males in the mixed treatment, including both, male-male and male-female (mating) interactions, fathered offspring of lower quality in both matings and for both lines (Fig. 3). Moreover, we find that male-male interactions (Irradiation × Intrasexual interactions: P_MCMC_ = 0.010) and mating (Irradiation × Intersexual interactions: P_MCMC_ = 0.088) decreased offspring quality after the short-term 3 h recovery period (Fig. 3A), albeit the latter effect was marginally non-significant. After the long-term recovery period (Fig. 3B), with males being held in isolation during the 24 h post irradiation resting period, the effect of male-male interactions was no longer detectable (Irradiation × Intrasexual interactions: P_MCMC_ = 0.844), while the effect of previous matings persisted (Irradiation × Intersexual interactions: P_MCMC_ = 0.030). The two lines differed overall in the observed reduction in offspring quality, but showed similar responses to the socio-sexual treatments (Fig. 3, for detailed statistics see: Supplementary Information S3 and S4).

**Figure 3:**
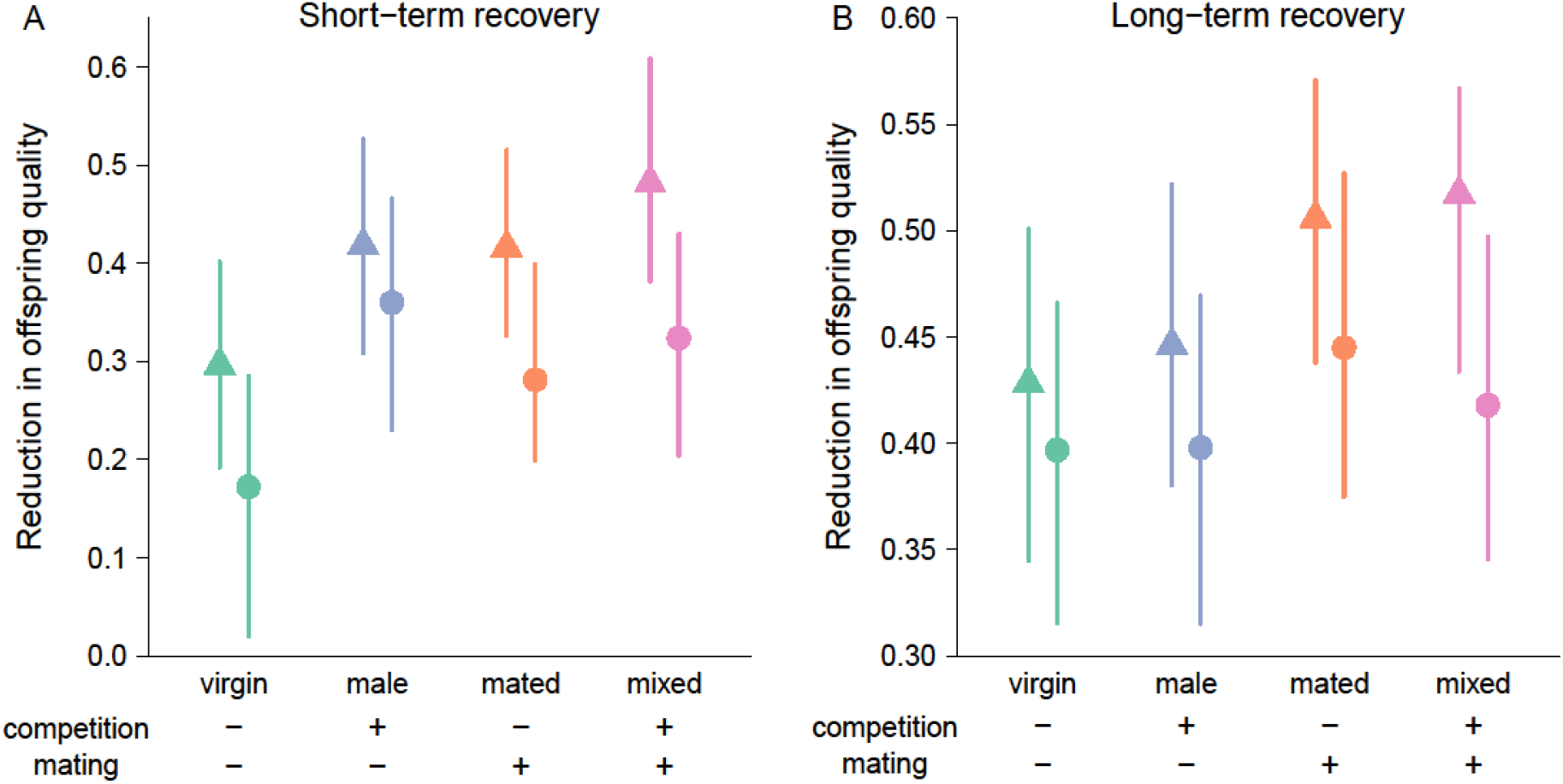
Reduction in offspring quality (1-[F2_IRRADIATED_/F2_CONTROL_]) after short-term (A) or long-term (B) recovery of males from two experimental evolution lines (S1: triangles, S3: circles) with an evolutionary history of intense sexual selection. Males were held in one of four different social environments before irradiation: solitary, without any competitors or mating partners (virgin, green symbols); without mating partners but with four male competitors (male, blue symbols); without competitors but with one female mating partner (mated, orange symbols); or with four male competitors and five female mating partners (mixed, pink symbols). Values are Bayesian posterior means and 95% credible intervals.

### Differential gene expression

To explore the molecular underpinnings of the reduction in germline maintenance, we took a subset of males from all treatment groups (line/irradiation/social environment) to analyze gene expression in the male reproductive tracts (Fig. 2C) after short term recovery. Analyzing all 12161 genes expressed in our data set, we found strong differences between the two experimental evolution lines (5910 DEGs, 49 %, Table 1) reflecting that these lines have been evolving separately for more than 50 generations. When looking at the effect of inter- and intrasexual interactions, most differences occurred in the environment that combined both types of interactions (mixed vs virgin, 3418 DEG, 28 %, Table 1), with the effect of mating (mated vs virgin, 2747 DEGs, 23 %, Table 1) contributing more than the effect of male-male competition (male vs virgin, 2 DEGs, < 1 %, Table 1). Irradiation resulted in only very few gene expression changes across both lines and all social environments (irradiated vs control, 18 DEGs, < 1 %, Table 1) and only one of those showed a larger than two-fold change (Table S1, Fig. 4B), which may in part be due to the timing of the measurements. Nevertheless, several of the DEGs are implicated in processes associated with germline maintenance and DNA repair. Among the up-regulated genes, three (*CALMAC_LOCUS18783*, *CALMAC_LOCUS9511* and *CALMAC_LOCUS2860*) code for proteins containing a MADF domain [70], which can also be found in the putative transcription factor s*tonewall* in *Drosophila melanogaster*, that is involved in female germline stem cell maintenance [71]. Also upregulated is the gene *CALMAC_LOCUS8201*, whose protein product contains a ULP protease domain [70]; in yeast, Ulp1 is involved in the sumoylation dynamics that play a critical role in the DNA damage response, specifically in the repair of double-strand breaks [72]. Among the down-regulated genes, *CALMAC_LOCUS9612* codes for a protein containing a BIR domain [70], which can also be found in the *D. melanogaster* apoptosis inhibitors Iap1 and Iap2 [73].

**Table 1:** Summary of differential gene expression analysis for the contrasts of interest, significance at 5 % False Discovery Rate

|  | Line<br>(MA3 – MA1) | Irradiation<br>(irrad. - control) | Male<br>(male - virgin) | Mated<br>(mated - virgin) | Mixed<br>(mixed - virgin) |
| --- | --- | --- | --- | --- | --- |
| down | 2867 | 8 | 2 | 1440 | 1720 |
| not sign. | 6251 | 12143 | 12159 | 9414 | 8743 |
| up | 3043 | 10 | 0 | 1307 | 1698 |

**Figure 4:**
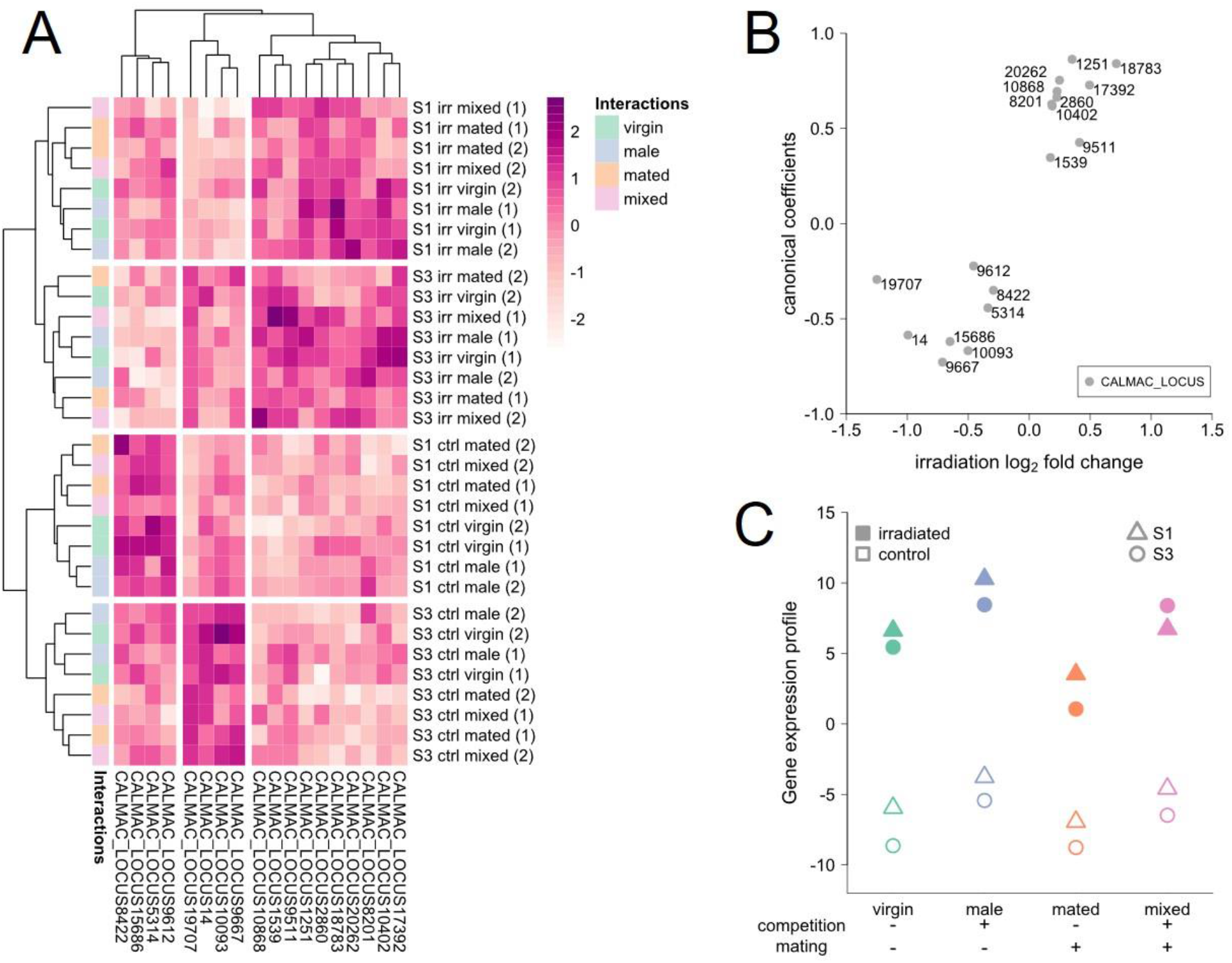
Gene expression in the reproductive tracts of S-males. (A) Heatmap of scaled normalized log2 expression of the 18 genes that responded to the irradiation treatment, expression is clearly separated between irradiation treatments (ctrl: control, irr: irradiated) and experimental evolution lines (S1, S3), within these blocks a separation between mated (orange and pink) and non-mated (green and blue) males can be observed. (B) Canonical coefficients of the 18 irradiation responsive genes that make up the canonical scores of each sample against their log2 fold change in response to irradiation. (C) Canonical scores separating control (open) and irradiated (closed) samples based on expression of the 18 irradiation responsive genes (triangles: S1; circles: S3).

Moreover, there was an overlap between genes responding to irradiation and to the social environments (specifically those treatments including intersexual interactions, Fig. S2). To explore effects of the social environment on irradiation responsive genes, we tested the 18 irradiation response candidate genes (Fig. 4A) in a MANOVA. Here, we took advantage of our full-factorial design and tested the interaction between intersexual interactions, intrasexual interactions and irradiation (Table 2). Both, inter- and intrasexual interactions significantly influenced overall expression of irradiation responsive genes independently (Table 2). Furthermore, intersexual interactions even significantly affected the irradiation response itself (Table 2).

**Table 2:** Test statistics of a multivariate analysis of variance with all 18 irradiation response genes as dependent variables

|  | **df** | **Pillai's trace** | **P** |
| --- | --- | --- | --- |
| **Line** | 1 | 0.98861 | **< 0.001** |
| **Irradiation** | 1 | 0.99632 | **< 0.001** |
| **Inter(sexual interactions)** | 1 | 0.98812 | **< 0.001** |
| **Intra(sexual interactions)** | 1 | 0.93388 | **0.032** |
| **Irradiation × Inter** | 1 | 0.96718 | **0.005** |
| **Irradiation × Intra** | 1 | 0.62005 | 0.852 |
| **Inter × Intra** | 1 | 0.70972 | 0.662 |
| **Irradiation × Inter × Intra** | 1 | 0.77902 | 0.452 |

To further explore this link, we first conducted a canonical discriminant analysis to find a linear combination of expression values of irradiation responsive genes that best separates the irradiation and control samples. To get the best representation of the irradiation effect while avoiding overfitting the data, we controlled for variation due to line, social environment and day and limited our interpretation to the first canonical axis. As expected, canonical coefficients for the 18 candidate genes roughly followed the log_2_ fold change induced by irradiation (Fig. 4B). This separation of irradiated from control samples revealed differences between the social environments (Fig. 4C). For example, males cohabitated with other males (intrasexual interactions) already have slightly more “irradiation-like” canonical scores in control groups, thus without any germline damage induced. This may indicate that they already have an increased need for germline maintenance due to elevated investment in germline replication or ejaculate composition in response to competition, or in response to direct stress induced by male rivals, as male-male competition is pronounced and carries considerable costs in this species [e.g. 74].

To determine whether the gene expression profiles of fathers from our treatment groups predicted the observed reduction in quality of offspring caused by the induced germline damage, we applied a canonical correlation analysis. Using the two lines and four socio-sexual treatments as units of replication, the gene expression canonical scores of control and irradiated F0 fathers (Fig. 4C) were entered as x variables, and the observed of reduction in offspring quality after the short and long recovery period (Fig. 3) as y variables. This thus resulted in 8 independent samples with 2 explanatory (gene expression) and 2 response (reduction in offspring quality) variables. Using F-approximations of Pillai-Bartlett’s trace both canonical dimensions taken together are significant (F = 7.10, df1 = 4, df2 = 10, P = 0.006) with the second dimension also being significant on its own (F = 6.37, df1 =1, df2 = 14, P = 0.024). Based on canonical dimension 1, more irradiation-like gene expression in control males was associated with greater reduction in the quality of offspring produced by the first mating following short term recovery (Table 3). The gene expression of control males, not themselves being exposed to irradiation, likely reflects the state of the germline as a result of the sociosexual interactions, suggesting that the observed correlation between gene expression in the reproductive tract of the fathers and the reduction in quality of their offspring was driven by male-male interactions (Fig. 5A). Canonical dimension 2 describes a correlation between the reduction in quality of offspring produced by the second mating following long-term recovery and the magnitude of the gene expression response to irradiation found in fathers (Table 3). This thus suggests that offspring quality is dependent on the capacity of fathers to modulate gene expression to deal with the induced damage, with stronger responses mitigating the consequences of germline damage. Here, the mating treatment, rather than male-male interactions, was the main driver of the correlation between germline gene expression and offspring quality (Fig. 5B).

**Table 3:** Canonical correlation coefficients

|  | Canonical coefficients |  | Standardized coefficients |  |
| --- | --- | --- | --- | --- |
|  | Dimension 1<br>(corr.: 0.92) | Dimension 2<br>(corr.: 0.79) | Dimension 1<br>(corr.: 0.92) | Dimension 2<br>(corr.: 0.79) |
| **Gene expression** |  |  |  |  |
| Control samples | -0.83 | -0.39 | -1.48 | -0.70 |
| Irradiated samples | 0.25 | 0.49 | 0.74 | 1.46 |
| **Reduction in offspring quality** |  |  |  |  |
| Short-term recovery | -11.71 | 10.38 | -1.14 | 1.01 |
| Long-term recovery | 4.27 | -33.16 | 0.19 | -1.51 |

**Figure 5:**
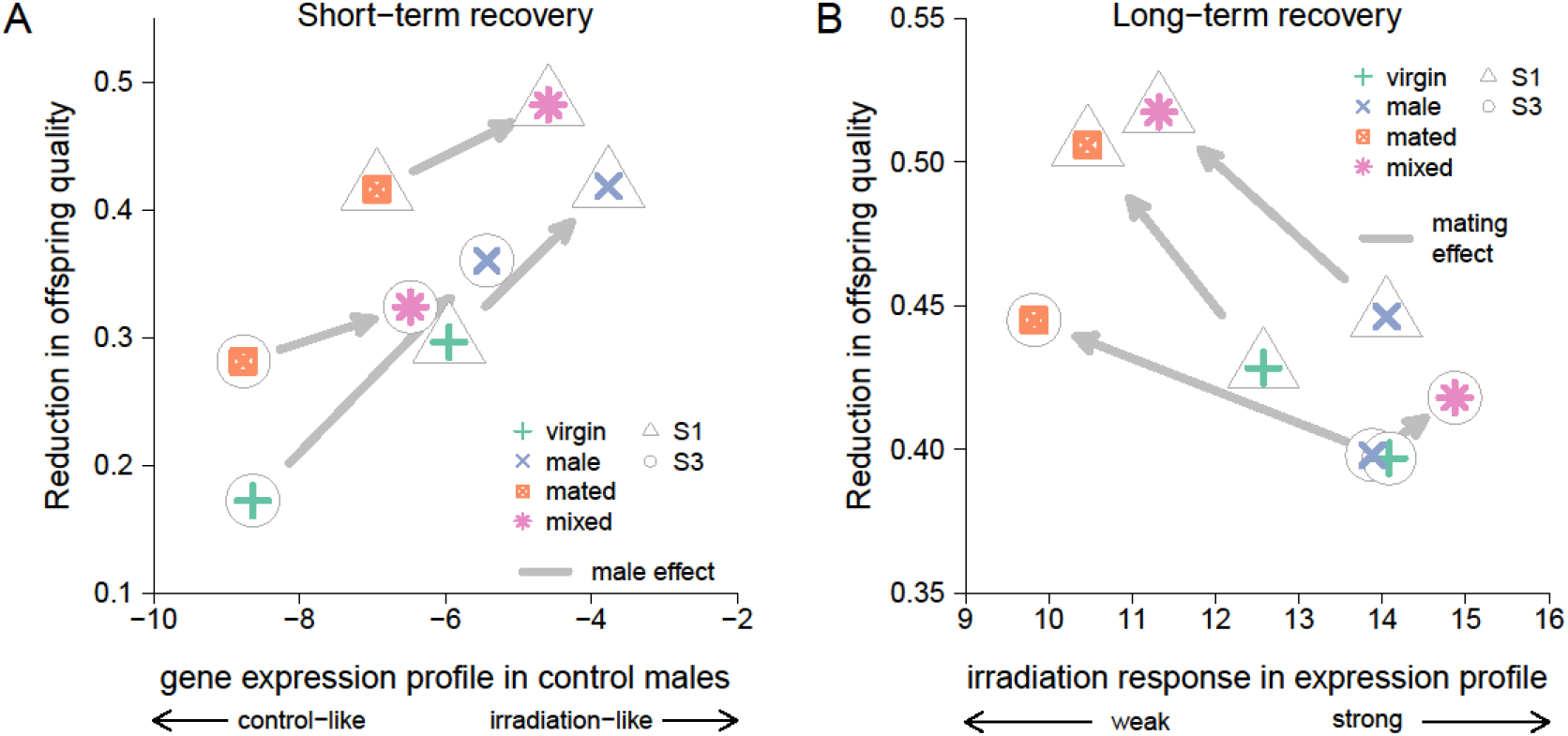
Relationship between gene expression profiles in fathers (as canonical scores, see Fig. 4) and the reduction in the quality of offspring fathered by irradiated males. (A) As indicated by the first dimension of the canonical correlation analysis, stronger reduction in offspring quality after a short-term recovery is associated with a more irradiation-like gene expression profile in control males (corresponding to baseline expression in socio-sexual treatments before irradiation), with male-male interactions leading to more irradiation-like gene expression profiles and stronger reduction in offspring quality. Arrows indicate the effect of adding males to the socio-sexual environment. (B) According to the second canonical dimension, larger gene expression response to irradiation (seen in unmated males), led to a smaller reduction in offspring quality. Arrows indicate the effect of adding females (and thus mating opportunities) to the socio-sexual environment.

#### Expression of irradiation responsive genes in experimental evolution lines

It has previously been established that males from the different selection regimes differ in their capacity to maintain their germline after induced damage (Fig. 2A and Baur and Berger (2020) [7]). We therefore compared the expression of the 18 irradiation responsive genes across all 8 experimental evolution lines. To this end, we analyzed available data that focused on the reproductive tissue. These data were RNA sequences from male and female abdomens from the experimental evolution lines, taken 24 h after a single mating, with females having access to beans and males being held in groups of five males during the 24 h period.

We first calculated canonical scores for males from all experimental evolution lines using the canonical coefficients from our previous analysis (Fig. 4B). While there was a tendency for S males to have a more irradiation-like gene expression profile than N and N+S males (Fig. 6B), we found no significant difference between the three experimental evolution regimes (ANOVA: F_2,5_ = 1.02, P = 0.425), which in part may reflect the low statistical power for the test, comparing regimes represented by only 2 or 3 replicate lines. In contrast, when analyzing sex differences across all 8 replicate lines, all but two genes showed a significant differential expression between males and females (Table S2). Genes upregulated due to irradiation tend to be female-biased and genes that are downregulated due to irradiation tend to be male-biased (Fisher’s exact test: P = 0.017; Figure 6A), indicating that females generally invest more heavily in germline maintenance than males do.

**Figure 6:**
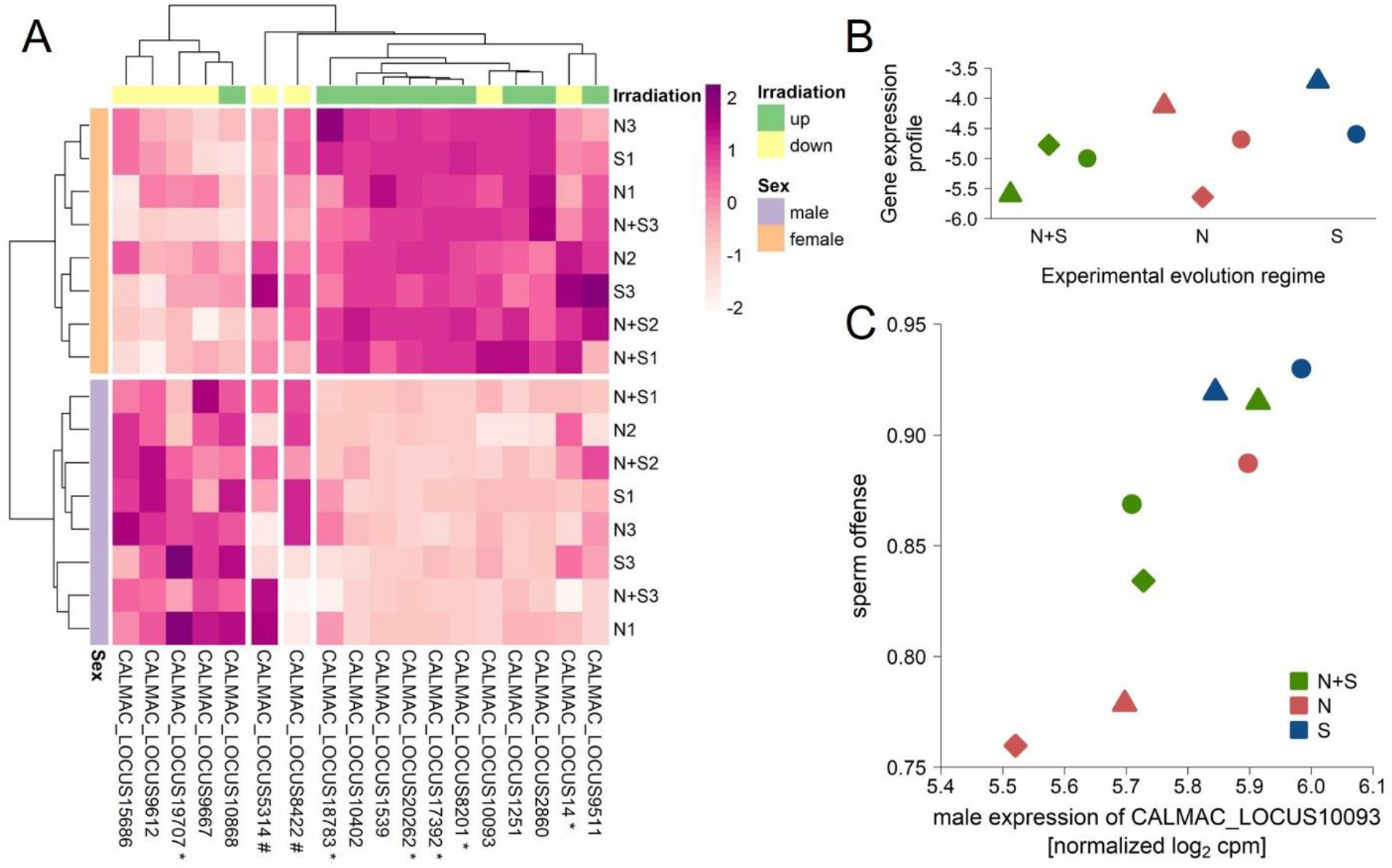
Expression of the 18 irradiation responsive genes in the 8 experimental evolution lines. (A) Heatmap of scaled normalized log2 expression values. Samples are separated by sex (females: orange; male: purple) and genes are separated by sex-bias, which roughly coincides with the direction of irradiation response (upregulated: green; downregulated: yellow). All but two genes (#, middle) show a significant differential expression between males and females, 6 genes show a significant sex-bias of at least twofold difference (*). Genes being upregulated in response to irradiation tend to be female-biased (right block), while genes being downregulated in response to irradiation tend to be male-biased (left block). (B) Scores (based on canonical coefficients used previously to separate control and irradiated samples) of male samples from the experimental evolution lines (24 h after a single mating). Higher scores indicate a more irradiation-like gene expression profile. (C) Positive correlation between normalized log_2_ expression of the irradiation responsive gene *CALMAC_LOCUS10093* (down-regulated in response to germline damage) and sperm offense (P2) ability of males from the experimental evolution lines. Experimental evolution regimes: N+S: natural and sexual selection; N: only natural selection; S :mainly sexual selection. Throughout the manuscript triangles refer to S1, N1, N+S1, diamonds refer to N2, N+S2 and circles refer to S3, N3, N+S3.

Our data are consistent with a trade-off between germline maintenance and investment in post-copulatory traits conferring advantages in sperm competition. Thus, we were interested in whether any of the irradiation responsive genes might be involved in sperm offense or defense. Therefore, we tested whether male expression of any of the 18 genes correlated with the estimated sperm competition ability (Fig. 1) in the 8 experimental evolution lines (Table S3 and S4). After p-value correction for multiple testing, one gene (*CALMAC_LOCUS10093*) retained a significant positive correlation with sperm offense (Fig. 6C). A higher expression of this gene is strongly statistically associated with a higher sperm offense success in *C. maculatus* males, while being downregulated in response to germline-damage. Since the predicted protein for this gene contains a *TFIIS N-terminal* domain [70], it is likely to be a transcription factor and thus may have a regulatory role in mediating the trade-off between investment into sperm competition and germline maintenance.

## Discussion

We hypothesized that male mutation rate and resulting offspring quality is governed by male strategies balancing the competing needs for post-copulatory reproductive success and germline maintenance. We therefore predicted that intense sexual selection coupled with the removal of constraints imposed by natural selection in the S regime would lead to the evolution of increased male reproductive competitiveness at the cost of germline maintenance.

We first confirmed a key expectation under this hypothesis by showing that S males had evolved increased post-copulatory reproductive success (Fig. 1). We then showed that previous findings, demonstrating that S males reduce germline maintenance when engaging in socio-sexual interactions [7], are repeatable and that male-male interactions (without mating) can be enough to elicit this response. We sequenced male reproductive tracts and identified 18 candidate genes that show differential expression in response to induced damage in the germline of S males. Many of those genes’ predicted protein products contain domains implicating their involvement in cellular maintenance and DNA repair. The 18 candidate genes also show significant expression changes across socio-sexual treatments and mating generally limited their damage-response, suggesting that these genes could be involved in a trade-off between germline replication and maintenance. We also found that the expression of these candidate genes in the reproductive tissue of fathers predicted the reduction in quality of their progeny brought about by the induced germline damage. Furthermore, we identified one gene whose expression was strongly positively correlated to sperm offense success but down-regulated in response to germline damage, suggesting that the gene could play a role in mediating the trade-off between post-copulatory reproductive success and offspring quality in *C. maculatus*. Our findings thus suggest that changes in the relative strengths of sexual and natural selection can lead to the evolution of phenotypic plasticity in the male reproductive tract with likely consequences for germline mutation rates and offspring quality.

Male reproductive investment in known to be responsive to female characteristics such as mating status [57] and to the presence of conspecific males [52, 58], as those cues can indicate the number of future mating opportunities and the level of sperm competition a male can expect [59]. Here, we found that germline maintenance was responsive to the presence of other males even in the absence of females and mating opportunities. This result is also in line with a recent study that found that intense male-male competition increases the fraction of sperm showing signs of DNA damage in zebrafish ejaculates [56]. While the need for investment into reproduction after mating is obvious as males need to replenish sperm and seminal fluid proteins, encounters with rival males do not result in an expenditure of those components, and thus may not necessarily stimulate further production of sperm and seminal fluid nor effectuate a reproduction-maintenance trade-off. Yet, encounters with conspecific males can serve as a signal for increased sexual competition that might warrant an increased investment into sperm and seminal fluid proteins that enhance post-copulatory fertilization success. Indeed, in *D. melanogaster*, male-male interactions change a male’s gene expression in the somatic (head/thorax) and reproductive (abdomen) tissue leading to the upregulation of several ejaculate component genes[75], indicating that males increase the production of seminal fluid proteins in response to perceived risk of sperm competition.

In addition to strategic allocation decisions in response to conspecific male competitors, it is also possible that the reduction in germline maintenance observed in our study was partly caused by the very costly nature of male-male interactions in *C. maculatus* [74]. In this scenario, manifestation of the trade-off between investment in reproduction (via engagement in reproductive competition) and germline maintenance could result from male-male competition depleting resources overall or shifting them away from the germline to the soma [9, 44]. This mechanism could also explain some discrepancies in the finer details of our results linking plastic responses in sperm competition success and germline maintenance. Most centrally, while we both here and previously [7] found that S males have evolved a plastic reduction in germline maintenance in response to social cues, we did not find that their success in sperm competition was improved by such cues (as expected in the trade-off scenario), nor that their response to these cues in terms of sperm competition success was different from that of the other regimes (although low statistical power may have played a role here). This, however, could be explained by male-male interaction being overall very detrimental to male condition[74], reducing sperm competition success. S males tend to engage more in such costly interactions than N, and possibly also N+S males [7]; therefore, because males were co-reared with males from their own regime in our experiments, it is possible that the detrimental effects of male cohabitation were most pronounced for S males in our assays of sperm competition success. Hence, even though there is differential allocation to sperm traits in response to social cues in S males compared to N and N+S males [7], any positive effects from this on sperm competition success may have been masked by stronger negative effects of male cohabitation in the S regime in our experiment.

Our gene expression data offer potential mechanistic insights into the allocation trade-off between sperm competition success and germline maintenance. In theory, the observed reductions in offspring quality fathered by males engaging in socio-sexual interactions could result from an increase in sperm production while keeping maintenance constant, rendering more replication errors unchecked per gamete. While this would not represent a functional allocation trade-off between maintenance and reproduction, it would still result in a trade-off between male success in sperm competition (assuming that success is dependent on sperm numbers) and gamete quality. Importantly, however, our gene expression data indicate that males engaging in mating interactions also have a decreased capacity to respond to DNA damage (Fig. 5B). Indeed, this result is in line with previous findings from these lines showing that reductions in offspring quality under inflated levels of germline damage is not a simple relationship of either sperm age or sperm production rate [7]. This study showed that socio-sexual treatment, the largest contributor to sperm replenishment and thus sperm age, had no general effect on offspring quality reduction, but rather only affected S males [7]. Furthermore, ejaculate investment patterns did not indicate a higher amount of sperm and seminal fluid being transferred by S males in a intrasexual competition setting compared to S-males in isolation, which makes sperm production rate an unlikely explanation for the observed offspring quality reduction [7].

Our gene expression data also suggest that *C. maculatus* males devote less resources into germline maintenance than their female counterparts. Given that sexual selection acts more strongly in males in most species [76, 77], including *C. maculatus* [78, 79], a trade-off between germline maintenance and postcopulatory reproductive success thus offers an explanation for the widely observed male mutation bias that goes beyond invoking sex differences in the numbers of germline replications as the sole driver of the phenomenon [7, 38, 40]. Data from other species are scarce, but there is some correlative comparative evidence to support a trade-off between germline mutation rate and post-copulatory reproductive investment. For example, testes mass – a possible adaptation to sperm competition - and substitution rates have been shown to covary in primates [80], and across bird taxa estimates of mutation rate have been shown to correlate with extra-pair paternity (as a measure of sperm competition intensity), but notably not with relative testes mass [39]. However, if these correlative patterns are indeed causal, and what their mechanistic explanation in that case may be, remains unknown.

In *C. maculatus*, males that are most successful in sperm offense sire daughters with lower lifetime reproductive success [81], which accords with the presence of substantial amounts of sexually antagonistic genetic variation for fitness in this species [82]. However, it is possible that this effect is partly mediated through reduced germline maintenance in successful males leading to lower genetic quality of their offspring. If so, a similar reduction in quality would also be expected for sons of successful males (which remains unconfirmed). This in turn may steer female choice away from these males in order to enhance the genetic quality of offspring [83]. However, systematic difference in male and female expression of genes involved in germline maintenance, as indicated here, and the capacity of females to repair damaged male sperm [6, 84], opens up for the possibility that females instead may devote large amounts of resources to maintenance of ejaculates and sperm deriving from males that are superior in sperm competition. Thus, male-female coevolution of germline maintenance may represent an important, yet understudied, aspect in the evolution of mate choice. More generally, the evolution of phenotypic plasticity in germline maintenance in response to sexual competition, as demonstrated here by experimental manipulation, might contribute to both, systematic differences in mutation rate between the sexes as well as among-male variation within species. Such variation has fundamental implications for theories of sexual selection, including reinforcement of mate choice and “good genes” processes, and for the maintenance of genetic variation in fitness related traits.

## Methods

### Beetles

The seed beetle *Callosobruchus maculatus* develops on the seeds of legumes. All beetles for the experiments were reared and kept on black-eyed beans (*Vigna unguiculata*) in constant climate chambers at 29 °C, 50 % relative humidity and a 12:12 L:D cycle. Where applicable, we used beetles from the ancestral *Lomé* population [see: 82, 85] as standardized mating partners and males from a black *C. maculatus* line as competitors [69]. Focal individuals came from 8 experimental evolution lines that originated from the *Lomé* population and are described in detail in [7, 68]. In short, those beetles evolved for > 50 generations under one of three experimental evolution regimes: N+S beetles evolved under polygamy with opportunities for natural (N) and sexual (S) selection to act, N beetles evolved under enforced monogamy with sexual competition between males removed and thus mainly natural (N) selection acting, S beetles evolved under polygamy but with a middle-class neighborhood breeding design applied to females weakening natural selection and leaving mainly sexual (S) selection to act. For the germline maintenance experiment, we focused on males from the two S lines, which were both mated to females from a third (polygamous) line to exclude any female derived and/or co-evolution effects. For the sperm competition experiment and the second gene expression dataset we included beetles from all 8 experimental evolution lines.

### Sperm competition

Males from all 8 experimental evolution lines were tested for the sperm competitiveness when being first (sperm defense, P1) or second (sperm offense, P2) to mate with a double mated female. To avoid potential confounding effects of females, we used females from the ancestral *Lomé* population that were mated to two males (observed single matings) 24 hours apart. Focal males were held in one of two social environments for approximately 24 h before their mating: solitary (single males in 30 mm dishes) or competition (in groups of five males in 90 mm dishes). The experiment was conducted twice. Within each of the two blocks, lines were separated into one of three sub-blocks, two sub-blocks contained one line from each of the three experimental evolution regimes and the last block contained the third replicate line of the N+S and N regime.

For sperm defense, females were mated to a focal male from one of the experimental evolution lines in a single observed mating and afterwards kept on beans for 24 h before their second mating. Beans were incubated for approximately 30 days and then frozen at −20 °C to determine 24 h offspring production elicited by the first mating. 24 h after the first mating all females were given the opportunity to mate with a black competitor male within 40 minutes. Successful mating pairs were separated and females were moved onto fresh beans for 48 h (N = 290, with a minimum of N = 26 per line). Beans were incubated for approximately 30 days and frozen at −20 °C, afterwards offspring was counted separated by black and wildtype fathers based on their coloration.

For sperm offense, males were consecutively mated five times to already mated females in one of the two blocks to determine the rate of decline in sperm competitiveness. Females were first mated to a black male in observed single matings and kept on beans for egg laying in groups of max. 25 for 24 h. Afterwards females were given the opportunity to mate with one focal male from the experimental evolution lines for 40 min. Successful pairs were separated, females from matings 1,3 and 5 were put on fresh beans for 48h to determine focal male sperm offense success (N = 784, with a minimum of N = 17 per line and mating). Beans were incubated for approximately 30 days and frozen at −20 °C. Afterwards, offspring was counted separated by black and wildtype fathers based on their morphology.

Statistical analyses and preparation of graphs were done in R 4.1.1 [86] using Bayesian Generalized Linear Models implemented within the package *MCMCglmm* [87]. Proportion of the focal (wildtype) father’s offspring was modelled with Binomial error distribution corrected for overdispersion. We included experimental evolution regime as well as its interaction with competition mode (sperm offense or defense), number of mating (three-level factor), and social environment (two-level factor) as fixed effects. To ease interpretation, non-significant interaction terms were removed stepwise. We included experimental evolution line crossed with competition mode, number of mating, and social environment as random terms. As additional random terms, we included experimental block, experimental evolution line, and male ID. For graphical presentation, line specific defense and offense success for a focal male’s first mating were calculated in separate models using package *MCMCglmm* [87] including block and social environment as random.

### Germline maintenance

#### Experimental assay

In order to measure germline maintenance capacity, we induced DNA-damage in adult males by exposing them to 25 Gray of gamma radiation (for 35 min at a dose rate of 0.72 Gray/min from a cesium-137 source). Gamma radiation causes double and single strand DNA breaks as well as increases the amount of reactive oxygen species in cells [34], which in turn can induce further DNA damage [34, 45]. While our treatment drastically increased germline damage, DNA breaks occur naturally during both recombination and chromatin remodeling during sperm development, and errors in the repair of those breaks give rise to point mutations [29, 34, 88]. The number of DNA lesions induced by a given dose of gamma radiation is surprisingly constant per DNA base pair [88], and thus differences in mutation rate are mainly caused by to the amount and type of repair molecules [34, 88, 89], which makes this assay ideal for measuring germline maintenance. Because most mutations are neutral or deleterious [90, 91], the amount of mutations transferred from parents to offspring can be approximated by the decline in offspring quality of parents that were challenged to deal with elevated levels of reactive oxygen species and DNA damage [6–8, 10, 15].

Assays were replicated on two consecutive days. Males and females (from the two S-lines) and females (from a third, polygamous line) were picked as virgins within 24 h after emergence. Females (from the third, polygamous line) were held in groups of ten in petri dishes (90 mm) until mating assays and males were immediately transferred to their respective socio-sexual environment using females from their own experimental evolution line where applicable. Socio-sexual treatments were set up by manipulating the presence of conspecific males and females in a full factorial design. Males were held in a 35 mm petri dish without any conspecifics or with a single virgin female, or in a 90 mm dish together with four conspecific males or with 4 conspecific males and an additional 5 virgin females. Males were held in their respective socio-sexual environment for approximately 24 h until shortly (< 1 h) before the irradiation treatment. Then, males were separated into individual 0.5 ml reaction tubes with a hole punched into the lid. Roughly half of the males then underwent a radiation treatment while the other half served as controls.

#### Offspring quality

Germline maintenance was assessed by measuring fitness effects of the induced germline damage in subsequent generations. Shortly after irradiation (1.5-3 h day 1, 2.5-4 h day 2) males were mated once to a single virgin female (0-48 h old) in a 60 mm petri dish on a heating plate set to 29 °C. Females were put on beans to lay eggs for 72 h and males remained in their individual petri dishes to renew their ejaculate, thus making sure that all males were challenged to deal with the competing tasks of both replicating and maintaining their germline. One day after irradiation (22-24 h day 1, 22-23 h day 2) males were again mated to a single virgin female (24-48 h old) in 60 mm dishes on a heating plate. Females were put on beans for 72 h to lay eggs and males were discarded. All beans were incubated at 29°C, 50 % r. h. and 12:12 L:D cycle in a climate chamber for 30 days to ensure that all viable offspring had emerged. Before offspring eclosion, beans were transferred to virgin chambers. Some of the offspring were used in the assay below. Remaining offspring were frozen at −20 °C and then counted to determine male offspring production from their first and second ejaculate.

To determine the reduction in quality of offspring fathered by irradiated males, we crossed the F_1_ offspring of each male with the F_1_ offspring of other males within the same treatment, experimental evolution line, and experimental day using a Middle-Class Neighbourhood breeding design (relaxing selection on the induced mutations). For the first ejaculate, we aimed at crossing one F_1_ male and one F_1_ female per male. For the second ejaculate, we aimed at crossing 3 F_1_ males and 3 F_1_ females, as we wished to focus on the recovery of the male germline. Pairs were kept on beans for their entire life and we incubated dishes for 33 days before freezing them at − 20°C to count F_2_ production. Counts of F_2_ adult offspring emerging from these irradiated lineages (n = 224) were compared to counts from corresponding control lineages (n = 163) to calculate reduction in offspring quality as: 1-[F2_IRRADIATED_/F2_CONTROL_]. Thus, we could explore phenotypic plasticity in germline maintenance in response to the socio-sexual treatments by comparing reduction in quality of offspring from grandfathers kept under the four treatments (Fig. 2B).

Again, we used Bayesian Generalized Linear Models implemented within the package *MCMCglmm* [87] in R 4.1.1 [86] for statistical analyses. Number of F_1_ and F_2_ offspring were modelled with Poisson error distribution corrected for overdispersion, for analysis of F_2_ offspring, dam and sire (IDs of the two grandfathers) were entered as a multiple-membership random term. Socio-sexual interactions were modelled as two two-level fixed effects (Inter- and Intrasexual interactions) testing for a significant interaction with irradiation treatment. We also added experimental evolution line and day as fixed effects to test for any differences between the two lines and the two assay days. To ease interpretation, non-significant interaction terms were removed. For graphical presentation, line specific means (and their 95% HPD interval) of the reduction in offspring quality were calculated per socio-sexual environment with Bayesian Generalized Linear Models and a Gaussian distribution. Similarly, those values were calculated for each of the two assay days separately for use in further analyses. Packages *Hmisc* [92] and *RColorBrewer* [93] were used to generate graphs.

#### Differential gene expression

For RNA extraction, beetles were snap frozen 2 h after irradiation treatment and stored at −80°C until dissections. During dissections, beetles and dissected tissues were kept on dry ice and afterwards stored at −80°C until RNA extraction. Males were dissected on ice in a droplet of PBS, the entire reproductive tract (Fig. 2C) was removed and the two large accessory gland pairs cut off. We decided to remove the two large accessory gland pairs in order to keep dissections consistent, as the large accessory glands easily detach and/or rupture during dissections. Afterwards the remaining tissue (aedeagus, ejaculatory bulb, 2 bilobed testes and 3 pairs of smaller accessory glands [ectadenial glands] [94]), was quickly rinsed in a fresh droplet of PBS and then transferred to a reaction tube on dry ice. We aimed to pool tissue from 10 males per sample, for two samples (1 mated irradiated line S3 and 1 mated control line S3) we only obtained tissue from nine males. Each sample consisted only of males from the same treatment, line and experimental day. This resulted in a total of 32 samples with 4 replicates per treatment (1 per day and line).

RNA was extracted with Qiagen RNeasy Mini Kit and on-column DNA digestion was performed with Qiagen RNase free DNase Kit. We followed the manufacturer’s instructions, beta-mercapto-Ethanol was added to the lysis buffer, tissue lysis was done with one stainless steel bead in a bead mill at 28 Hz for 90 s. Two samples underwent an additional clean-up using the Qiagen RNeasy Mini Kit. RNA concentration and purity were assessed with NanoDrop and additional quality controls were performed at the sequencing facility. Samples were sequenced at the SNP&SEQ Technology Platform in Uppsala. Libraries were multiplexed and sequenced as stranded paired-end 50 bp reads in 2 lanes of a NovaSeq SP flow cell resulting in roughly 11M-26M reads per sample.

Raw reads were inspected with *FastQC* [95] and quality information summarized with *MultiQC* [96]. We then mapped all reads to the *C. maculatus* genome (GCA_900659725.1; ASM90065972v1) [97] with *TopHat 2.1.1* [98] allowing for up to two mismatches per read and including strand information. We only kept reads where both mates successfully mapped to the *C. maculatus* genome. Those reads were then counted per gene using *HTSeq* [99] with default settings for stranded libraries. Statistical analyses were done in R 4.1.1 [86], with packages *edgeR* [100] and *limma* [101]. Libraries from the two lanes were merged into one sample. Genes that were not at least expressed as 1 count per million (cpm) in at least 2 samples were excluded from the analysis resulting in a total of 12161 genes being analyzed. Counts were normalized with the ‘Trimmed Mean of M-values’ method and normalized log_2_ cpm values were analyzed in linear models within *limma* [101]. Experimental evolution line and experimental day were added as additive terms to control for variance between lines and days. Socio-sexual environment was entered as a 4-level factor and irradiation treatment as a two-level factor. Due to the low number of genes responding to the irradiation treatment, we lacked statistical power to analyze the interaction between social environment and irradiation with the full set of genes. Therefore, the interaction was removed from the model and we analyzed the interaction in a separate model considering only genes that responded to the irradiation treatment. We used information available on *UniProt* [70] accessed on 10.03.2022 to gain insight on the potential function of some of the genes found to be differentially expressed.

For further analyses we always used normalized log_2_ cpm values. Using the 18 irradiation responsive genes, we ran a multivariate Anova. Additionally, we ran a linear discriminant analysis on gene expression in the 18 irradiation responsive genes to find a linear combination of the expression of these genes that best separates irradiated from control samples. To this end we controlled for variation arising through differences in irradiation response between lines, socio-sexual environments or experimental days using package *candisc* [102]. To avoid overfitting the data, we calculated canonical scores of the 32 samples with the first canonical axis only. For further analyses and graphical representation, we used mean canonical scores across the two experimental days. To estimate how well differences in gene expression correspond to differences in reduction in offspring quality due to germline damage, we conducted a Canonical Correlation Analysis. We added canonical scores of irradiated and control samples (averaged across the two experimental days) as two x variables and reduction in offspring quality after a short- and long-term recovery period as two y variables to the Canonical Correlation Analysis implemented in the package *CCA* [103]. For graphical presentation, heatmaps were constructed on scaled normalized log_2_ cpm values using hierarchical clustering and Manhattan distance metrics in *pheatmap* [104], and additional packages *VennDiagram* [105] and RColorBrewer [93] were used.

### Expression of irradiation responsive genes in experimental evolution lines

To analyze the expression of irradiation responsive genes in the 8 lines from all 3 experimental evolution regimes, we made use of an existing data set designed to study the evolution of sex-biased gene expression under these selection regimes. Before collecting individuals for sequencing, all experimental evolution lines underwent 3 generations of common garden rearing (i.e., a polygamous mating setting). Males and females from the experimental evolution lines were mated once (observed) to a standardized mating partner from the *Lomé* population on heating plates set to 29 °C. Matings were separated into 4 blocks and in each block, we set up 6 mating pairs per line and sex. Beetles from the first 5 successful matings per line and sex were separated after the end of the mating, focal females were kept singly on beans for 24 h and focal males were held together in a 90 mm dish (in groups of 5 individuals) for 24 h.

After 24 h, males and females were flash frozen in liquid nitrogen and kept at −80 °C until sample preparation. Since we were interested in the reproductive tissues, we only sampled the abdomen of males and females. To that end, we separated the abdomen from the rest of the body on ice, while storing samples on dry ice during preparation. We pooled 12 abdomen per group balanced over the four blocks (except for the male sample from line N+S2, which only contained 10 abdomen, block information on the two lost abdomen is not available). After dissection, samples were stored again at −80 °C until RNA extraction.

RNA was extracted with Qiagen RNeasy Mini Kit and on-column DNA digestion was performed with Qiagen RNase free DNase Kit. We followed the manufacturer’s instructions, beta-mercapto-Ethanol was added to the lysis buffer and tissue lysis was done with two stainless steel beads in a bead mill at 28 Hz for 90 s in 700 μl lysis buffer. After centrifugation the entire supernatant was transferred to a fresh tube, mixed quickly and 350 μl went onto the extraction column while the rest was discarded to avoid overloading the column. RNA was eluted 2 times in 50 μl of RNase free water each. RNA concentration and purity were assessed with NanoDrop, gel electrophoresis and Qbit, additional quality controls were performed at the sequencing facility. Samples were sequenced at the SNP&SEQ Technology Platform in Uppsala. Libraries were multiplexed and sequenced as stranded paired-end 150 bp reads in 1 lane of a NovaSeq S4 flow cell resulting in roughly 24M-56M reads per sample.

Gene counts were obtained and analyzed as in the main experiment with the exception of an additional quality and adapter trimming step with *Trimmomatic* [106] and allowing for up to eight mismatches per read during mapping due to the higher read length. Normalized log_2_ cpm counts of all 12874 retained genes were analyzed in a linear model within *limma* [101]. We constructed an additive model with sex (2-level factor) and experimental evolution regime (3-level factor) as explanatory variables. Sex bias was estimated across all experimental evolution regimes (male - female) and p-values were corrected with Benjamini-Hochberg method using a 5% FDR cut-off. We then extracted normalized log_2_ cpm values of the 18 irradiation responsive genes for all samples for further analysis. Using these values, we predicted canonical scores for males from all experimental evolution lines based on the linear coefficients from the previous analysis.

## Declarations

### Ethics approval and consent to participate

Not applicable.

### Consent for publication

Not applicable.

### Authors’ Contributions

MK, JB and DB designed the study, MK and JB conducted the research, MK and DB analysed the data and drafted the manuscript. All authors contributed to the final version of the manuscript.

## Acknowledgements

We thank P. E. Eady for sharing the *C. maculatus* black line with us and J. Liljestrand-Rönn for help in the lab.

