## Supplementary Information for "Increased male investment in sperm competition results in reduced maintenance of gametes"

#### S1: Results on fertility reduction

Reduced offspring production in the F1 generation is mainly driven by two mechanisms. Mutations carried by the male's sperm that fertilizes a female's ova carry dominant lethal mutations that lead to the fertilized egg not developing into an adult. Furthermore, radiation effects on male sperm morphology and seminal fluid proteins may diminish male fertilization ability and his ability to elicit mating induced fecundity boost in females (paternal effects). For the analysis, we excluded males that produced no offspring (short-term recovery: 2 out of 168 in controls and 8 out of 233 in irradiated; long-term recovery: 0 out of 405), as we assume that these are rather caused by a failed mating than the radiation effect. Irradiation had a strong negative effect on male fertility after both, long- and short-term recovery, with up ~ 80 % reduction in fertility (Fig. S1). Line S3 was significantly less affected by radiation than line S1 but there were no other significant differences between the two lines (for detailed statistics see further below). Overall, the interaction between inter- and intrasexual interactions did not influence fertility reduction and we could only observe a significant effect of mating on fertility reduction after short-term recovery ($P_{MCMC} = 0.03$) with mating opportunities prior to irradiation increasing the negative effect of radiation on fertility.

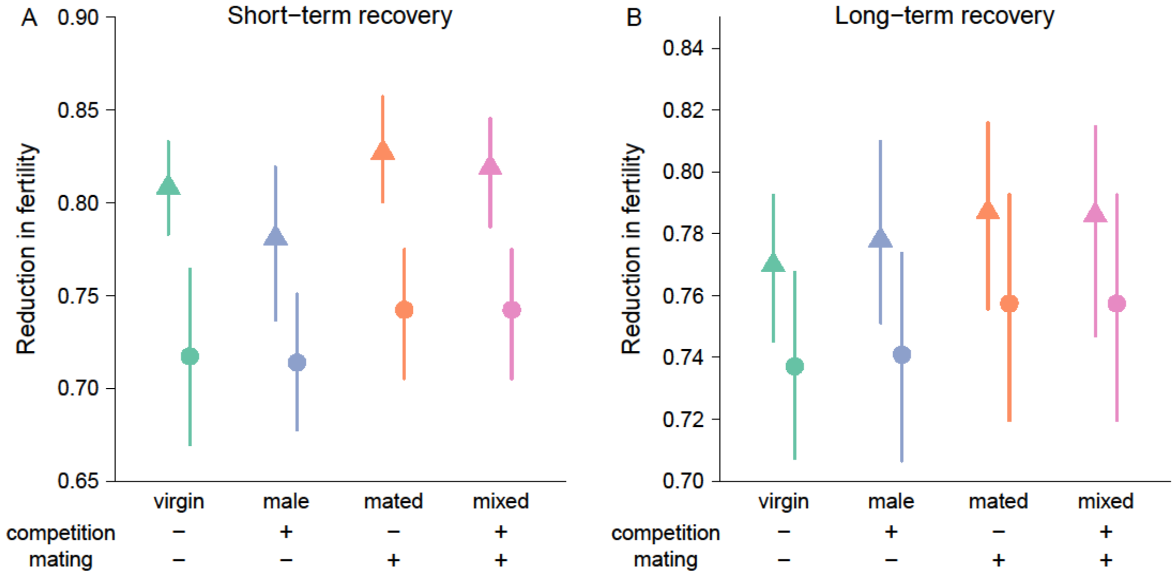

Figure S1: Irradiation induced fertility reduction (1-[offspring irradiated/offspring control]) after a short- (A) and long- (B) term recovery period for lines S1 (triangles) and S3 (circles). Males have been in one of four different socio-sexual environments prior to irradiation manipulating the presence of conspecific males (virgin and mated without other males, male and mixed with male competitors) and females (virgin and male without mating opportunities, mated and mixed with matin opportunities).

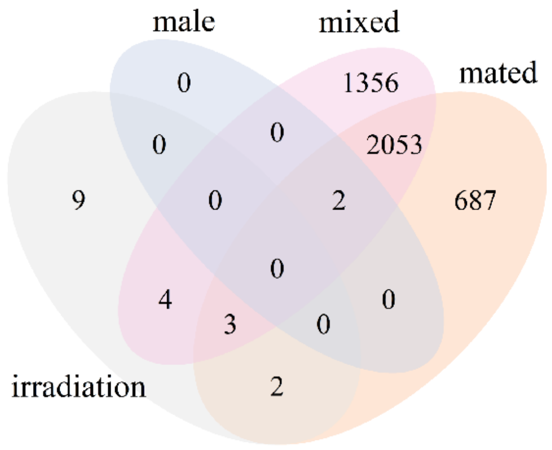

Figure S2: Venn Diagram of differentially expressed genes (5 % false discovery rate) of the main contrasts (socio-sexual treatments are all compared to virgin males).

Table S1: Test statistics and $log_2$-fold change (irradiated - control) of the 18 genes being significantly differentially expressed between irradiated and control samples (at 5 % false discovery rate).

| **Gene** | **$Log_2$-fold change** | **Average expression** | **t** | **P-Value** | **BH adjusted P-Value** | **B** |
|---|---|---|---|---|---|---|
| *CALMAC_LOCUS19707* | -1.250 | 0.541 | -5.933 | 1.84E-06 | 0.003 | 4.840 |
| *CALMAC_LOCUS14* | -0.994 | 2.436 | -7.370 | 3.80E-08 | < 0.001 | 8.104 |
| *CALMAC_LOCUS18783* | 0.715 | 0.827 | 6.657 | 2.54E-07 | 0.001 | 5.280 |
| *CALMAC_LOCUS9667* | -0.711 | 3.069 | -5.804 | 2.63E-06 | 0.004 | 4.643 |
| *CALMAC_LOCUS15686* | -0.650 | 3.636 | -6.030 | 1.41E-06 | 0.003 | 5.288 |
| *CALMAC_LOCUS10093* | -0.502 | 5.119 | -5.522 | 5.76E-06 | 0.006 | 3.970 |
| *CALMAC_LOCUS17392* | 0.495 | 3.819 | 6.690 | 2.32E-07 | 0.001 | 6.980 |
| *CALMAC_LOCUS9612* | -0.456 | 2.424 | -4.796 | 4.37E-05 | 0.031 | 2.046 |
| *CALMAC_LOCUS9511* | 0.413 | 1.695 | 5.214 | 1.36E-05 | 0.012 | 2.862 |
| *CALMAC_LOCUS1251* | 0.352 | 6.058 | 8.944 | 7.12E-10 | < 0.001 | 12.569 |
| *CALMAC_LOCUS5314* | -0.338 | 2.601 | -5.572 | 5.01E-06 | 0.006 | 3.981 |
| *CALMAC_LOCUS8422* | -0.294 | 2.511 | -4.843 | 3.83E-05 | 0.029 | 2.171 |
| *CALMAC_LOCUS20262* | 0.248 | 3.280 | 5.246 | 1.24E-05 | 0.012 | 3.259 |
| *CALMAC_LOCUS10868* | 0.228 | 6.723 | 6.685 | 2.36E-07 | 0.001 | 7.029 |
| *CALMAC_LOCUS2860* | 0.227 | 4.565 | 4.871 | 3.54E-05 | 0.029 | 2.258 |
| *CALMAC_LOCUS10402* | 0.188 | 7.123 | 5.561 | 5.17E-06 | 0.006 | 4.062 |
| *CALMAC_LOCUS8201* | 0.185 | 5.119 | 5.422 | 7.61E-06 | 0.008 | 3.702 |
| *CALMAC_LOCUS1539* | 0.174 | 5.122 | 4.778 | 4.60E-05 | 0.031 | 1.980 |

Table S2: Sex bias (male - female) in expression of the 18 irradiation responsive genes. Table represents a subset taken from the overall sex-bias analysis in all 12874 analyzed genes.

| **Gene** | **$Log_2$-fold change** | **Average expression** | **t** | **P-Value** | **BH-adjusted P-Value** | **B** |
|---|---|---|---|---|---|---|
| *CALMAC_LOCUS10093* | -0.700 | 6.141 | -11.883 | 1.34E-08 | 5.01E-08 | 9.298 |
| *CALMAC_LOCUS10402* | -0.906 | 7.672 | -15.586 | 4.13E-10 | 2.49E-09 | 12.887 |
| *CALMAC_LOCUS10868* | 0.467 | 6.849 | 8.823 | 5.06E-07 | 1.25E-06 | 5.373 |
| *CALMAC_LOCUS1251* | -0.549 | 6.519 | -10.298 | 7.89E-08 | 2.39E-07 | 7.376 |
| *CALMAC_LOCUS14* | -1.433 | 4.042 | -3.148 | 7.28E-03 | 9.73E-03 | -4.073 |
| *CALMAC_LOCUS1539* | -0.956 | 5.860 | -16.988 | 1.34E-10 | 9.85E-10 | 14.222 |
| *CALMAC_LOCUS15686* | 0.555 | 4.788 | 3.820 | 1.94E-03 | 2.77E-03 | -2.991 |
| *CALMAC_LOCUS17392* | -1.828 | 5.991 | -33.818 | 1.33E-14 | 9.54E-13 | 23.904 |
| *CALMAC_LOCUS18783* | -1.116 | 0.557 | -4.277 | 8.03E-04 | 1.19E-03 | -0.993 |
| *CALMAC_LOCUS19707* | 1.599 | -0.794 | 3.133 | 7.50E-03 | 1.00E-02 | -2.976 |
| *CALMAC_LOCUS20262* | -1.957 | 4.492 | -21.700 | 5.24E-12 | 7.33E-11 | 17.882 |
| *CALMAC_LOCUS2860* | -0.901 | 5.181 | -8.235 | 1.13E-06 | 2.60E-06 | 4.738 |
| *CALMAC_LOCUS5314* | -0.083 | 3.532 | -0.785 | 4.46E-01 | 4.75E-01 | -7.686 |
| *CALMAC_LOCUS8201* | -1.262 | 5.952 | -26.018 | 4.61E-13 | 1.17E-11 | 20.205 |
| *CALMAC_LOCUS8422* | -0.059 | 2.672 | -0.537 | 6.00E-01 | 6.26E-01 | -7.636 |
| *CALMAC_LOCUS9511* | -0.368 | 1.628 | -2.422 | 2.99E-02 | 3.72E-02 | -4.888 |
| *CALMAC_LOCUS9612* | 0.669 | 2.806 | 6.293 | 2.18E-05 | 3.98E-05 | 2.125 |
| *CALMAC_LOCUS9667* | 0.719 | 4.503 | 5.137 | 1.62E-04 | 2.61E-04 | -0.366 |

Table S3: Pearson's correlation coefficients and test statistics of individual correlation analyses between sperm defense success (P1) and expression of irradiation responsive genes in experimental evolution lines. Multiple testing correction of P-Values was done with Benjamini-Hochberg (BH) method.

| **Gene** | **Pearson's *r*** | **$t_6$** | **P-Value** | **BH-adjusted P-Value** |
|---|---|---|---|---|
| *CALMAC_LOCUS19707* | 0.08 | 0.209 | 0.842 | 0.947 |
| *CALMAC_LOCUS14* | 0.59 | 1.808 | 0.121 | 0.543 |
| *CALMAC_LOCUS18783* | -0.09 | -0.226 | 0.829 | 0.947 |
| *CALMAC_LOCUS9667* | -0.02 | -0.046 | 0.965 | 0.965 |
| *CALMAC_LOCUS15686* | -0.43 | -1.177 | 0.284 | 0.639 |
| *CALMAC_LOCUS10093* | -0.02 | -0.045 | 0.965 | 0.965 |
| *CALMAC_LOCUS17392* | 0.73 | 2.644 | 0.038 | 0.345 |
| *CALMAC_LOCUS9612* | -0.30 | -0.765 | 0.473 | 0.774 |
| *CALMAC_LOCUS9511* | -0.49 | -1.366 | 0.221 | 0.568 |
| *CALMAC_LOCUS1251* | -0.54 | -1.567 | 0.168 | 0.568 |
| *CALMAC_LOCUS5314* | -0.38 | -1.007 | 0.353 | 0.663 |
| *CALMAC_LOCUS8422* | 0.12 | 0.307 | 0.769 | 0.947 |
| *CALMAC_LOCUS20262* | 0.18 | 0.455 | 0.665 | 0.947 |
| *CALMAC_LOCUS10868* | 0.77 | 2.984 | 0.025 | 0.345 |
| *CALMAC_LOCUS2860* | -0.15 | -0.362 | 0.730 | 0.947 |
| *CALMAC_LOCUS10402* | -0.62 | -1.949 | 0.099 | 0.543 |
| *CALMAC_LOCUS8201* | 0.49 | 1.391 | 0.213 | 0.568 |
| *CALMAC_LOCUS1539* | -0.37 | -0.973 | 0.368 | 0.663 |

Table S4: Pearson's correlation coefficients and test statistics of individual correlation analyses between sperm offense success (P2) and expression of irradiation responsive genes in experimental evolution lines. Multiple testing correction of P-Values was done with Benjamini-Hochberg (BH) method.

| **Gene** | **Pearson's *r*** | **$t_6$** | **P-Value** | **BH-adjusted P-Value** |
|---|---|---|---|---|
| *CALMAC_LOCUS19707* | 0.16 | 0.398 | 0.704 | 0.906 |
| *CALMAC_LOCUS14* | -0.23 | -0.577 | 0.585 | 0.906 |
| *CALMAC_LOCUS18783* | -0.16 | -0.407 | 0.698 | 0.906 |
| *CALMAC_LOCUS9667* | -0.10 | -0.248 | 0.813 | 0.912 |
| *CALMAC_LOCUS15686* | -0.22 | -0.552 | 0.601 | 0.906 |
| ***CALMAC_LOCUS10093*** | **0.90** | **5.196** | **0.002** | **0.036** |
| *CALMAC_LOCUS17392* | 0.21 | 0.535 | 0.612 | 0.906 |
| *CALMAC_LOCUS9612* | 0.19 | 0.483 | 0.647 | 0.906 |
| *CALMAC_LOCUS9511* | 0.37 | 0.987 | 0.362 | 0.906 |
| *CALMAC_LOCUS1251* | 0.18 | 0.442 | 0.674 | 0.906 |
| *CALMAC_LOCUS5314* | -0.23 | -0.57 | 0.589 | 0.906 |
| *CALMAC_LOCUS8422* | 0.12 | 0.294 | 0.779 | 0.912 |
| *CALMAC_LOCUS20262* | -0.24 | -0.593 | 0.575 | 0.906 |
| *CALMAC_LOCUS10868* | 0.00 | 0.009 | 0.993 | 0.993 |
| *CALMAC_LOCUS2860* | 0.54 | 1.591 | 0.163 | 0.906 |
| *CALMAC_LOCUS10402* | -0.27 | -0.696 | 0.513 | 0.906 |
| *CALMAC_LOCUS8201* | 0.57 | 1.685 | 0.143 | 0.906 |
| *CALMAC_LOCUS1539* | -0.07 | -0.183 | 0.861 | 0.912 |

### S2: Sperm competition, statistics summary

#### **Final reduced model**

Iterations = 200001:5195001
Thinning interval = 5000
Sample size = 1000
DIC: 28121.45

G-structure:

| | post.mean | l-95% CI | u-95% CI | eff.samp |
|---|---|---|---|---|
| Line | 0.01414 | 1.36e-07 | 0.06878 | 895.2 |
| Line:mating | 0.05134 | 2.316e-07 | 0.273 | 1000 |
| Line:Paternity | 0.02705 | 1.921e-07 | 0.1521 | 1000 |
| Line:Environment | 0.01124 | 1.377e-07 | 0.06353 | 1000 |
| Block | 0.1796 | 8.913e-08 | 0.6223 | 1000 |
| ID | 0.08455 | 1.817e-07 | 0.4673 | 592.8 |

R-structure: ~units

| | post.mean | l-95% CI | u-95% CI | eff.samp |
|---|---|---|---|---|
| units | 6.426 | 5.612 | 7.317 | 878.4 |

Location effects: cbind(Wt, Black) ~ Regime + Paternity + mating + Environment

| | post.mean | l-95% CI | u-95% CI | eff.samp | pMCMC |
|---|---|---|---|---|---|
| (Intercept)[a] | -0.54250 | -1.18542 | 0.28785 | 1195 | 0.122 |
| **RegimeMono** | **-0.77671** | **-1.37909** | **-0.10157** | **1271** | **0.018** |
| **RegimePoly** | **-0.51936** | **-1.09179** | **0.12260** | **1148** | **0.078** |
| PaternityP2 | 2.64973 | 2.21965 | 3.10592 | 1000 | <0.001 |
| matingM3 | -0.58138 | -1.16437 | -0.08821 | 1000 | 0.048 |
| matingM5 | -1.93049 | -2.44442 | -1.25492 | 1000 | <0.001 |
| Environmentsolitary | 0.27576 | -0.08476 | 0.61090 | 1000 | 0.132 |

[a]The intercept represents sperm defense (P1) in the first mating in S-males after being held in competition (4 other males) 24 h prior to the sperm competition assay.

### S3: Reduction in offspring quality after short-term recovery period, statistics summary

#### Final reduced model

Iterations = 200001:5195001
Thinning interval = 5000
Sample size = 1000
DIC: 3087.496

G-structure: ~mm(Dam_n + Sire_n)

| | post.mean | l-95% CI | u-95% CI | eff.samp |
|---|---|---|---|---|
| Dam_n+Sire_n | 0.0009809 | 1.56e-07 | 0.003903 | 384.8 |

R-structure: ~idh(Irradiation:OtherMales:Mating):units

| | post.mean | l-95% CI | u-95% CI | eff.samp |
|---|---|---|---|---|
| Irradiationctrl:OtherMalesNoMales:Matingmated.units | 0.0006591 | 1.394e-07 | 0.003497 | 1000 |
| Irradiationirr:OtherMalesNoMales:Matingmated.units | 0.1381204 | 7.957e-02 | 0.201832 | 1000 |
| Irradiationctrl:OtherMalesOtherMales:Matingmated.units | 0.0102144 | 3.576e-07 | 0.020712 | 607 |
| Irradiationirr:OtherMalesOtherMales:Matingmated.units | 0.3371629 | 2.073e-01 | 0.502774 | 1000 |
| Irradiationctrl:OtherMalesNoMales:Matingvirgin.units | 0.0114436 | 4.548e-07 | 0.023814 | 1000 |
| Irradiationirr:OtherMalesNoMales:Matingvirgin.units | 0.2051549 | 1.248e-01 | 0.294872 | 1512 |
| Irradiationctrl:OtherMalesOtherMales:Matingvirgin.units | 0.0201414 | 5.251e-03 | 0.034927 | 1000 |
| Irradiationirr:OtherMalesOtherMales:Matingvirgin.units | 0.2757921 | 1.662e-01 | 0.397390 | 1000 |

Location effects: Offspring ~ Irradiation * OtherMales + Irradiation * Mating + Irradiation * Line

| | post.mean | l-95% CI | u-95% CI | eff.samp | pMCMC |
|---|---|---|---|---|---|
| (Intercept)[a] | 4.643279 | 4.596021 | 4.687729 | 1000 | <0.001 |
| Irradiationirr | -0.591616 | -0.718332 | -0.475799 | 1105 | <0.001 |
| OtherMalesOtherMales | 0.001035 | -0.041375 | 0.051118 | 1000 | 0.988 |
| Matingvirgin | -0.017207 | -0.069179 | 0.026461 | 1000 | 0.488 |
| LineMA3 | -0.005548 | -0.048280 | 0.042137 | 1000 | 0.794 |
| **Irradiationirr:OtherMalesOtherMales** | **-0.204506** | **-0.329832** | **-0.051711** | **1012** | **0.010** |
| **Irradiationirr:Matingvirgin** | **0.121672** | **-0.023893** | **0.250948** | **1000** | **0.088** |
| Irradiationirr:LineMA3 | 0.218059 | 0.083391 | 0.363522 | 1000 | 0.002 |

[a]The intercept represents the control treatment with no other males but a female mating partner in the socio-sexual environment (group “mated”) in males from line S1.

#### S4: Reduction in offspring quality after long-term recovery period, statistics summary

##### Final reduced model

Iterations = 200001:5195001
Thinning interval = 5000
Sample size = 1000
DIC: 9599.919

G-structure: ~mm(Dam_n + Sire_n)

| | post.mean | l-95% CI | u-95% CI | eff.samp |
|---|---|---|---|---|
| Dam_n+Sire_n | 0.00044 | 1.404e-07 | 0.002097 | 1000 |

R-structure: ~idh(Irradiation:OtherMales:Mating):units

| | post.mean | l-95% CI | u-95% CI | eff.samp |
|---|---|---|---|---|
| Irradiationctrl:OtherMalesNoMales:Matingmated.units | 0.02293 | 0.01433 | 0.03216 | 1000.0 |
| Irradiationirr:OtherMalesNoMales:Matingmated.units | 0.60516 | 0.45373 | 0.75083 | 1000.0 |
| Irradiationctrl:OtherMalesOtherMales:Matingmated.units | 0.05947 | 0.03693 | 0.08438 | 1144.0 |
| Irradiationirr:OtherMalesOtherMales:Matingmated.units | 0.56492 | 0.42483 | 0.71519 | 1000.0 |
| Irradiationctrl:OtherMalesNoMales:Matingvirgin.units | 0.06223 | 0.03912 | 0.08860 | 1000.0 |
| Irradiationirr:OtherMalesNoMales:Matingvirgin.units | 0.55783 | 0.43234 | 0.70548 | 960.5 |
| Irradiationctrl:OtherMalesOtherMales:Matingvirgin.units | 0.10257 | 0.06552 | 0.13963 | 1000.0 |
| Irradiationirr:OtherMalesOtherMales:Matingvirgin.units | 0.50336 | 0.38070 | 0.63391 | 1000.0 |

Location effects: Offspring ~ Irradiation * Mating + Irradiation * OtherMales + Irradiation * Line + Irradiation * Day

| | post.mean | l-95% CI | u-95% CI | eff.samp | pMCMC |
|---|---|---|---|---|---|
| (Intercept) [a] | 4.6214789 | 4.5737013 | 4.6665965 | 1096.5 | <0.001 |
| Irradiationirr | -0.8551001 | -0.9992089 | -0.7289456 | 1124.5 | <0.001 |
| Matingvirgin | -0.0358189 | -0.0849920 | 0.0084509 | 1000.0 | 0.130 |
| OtherMalesOtherMales | 0.0018353 | -0.0477967 | 0.0441239 | 964.0 | 0.930 |
| LineMA3 | 0.0290843 | -0.0119659 | 0.0720544 | 1000.0 | 0.180 |
| Day2 | 0.0010393 | -0.0468999 | 0.0451672 | 913.1 | 0.990 |
| **Irradiationirr:Matingvirgin** | **0.1323434** | **0.0207648** | **0.2667268** | **1000.0** | **0.030** |
| **Irradiationirr:OtherMalesOtherMales** | **0.0116687** | **-0.1035681** | **0.1373192** | **1000.0** | **0.844** |
| Irradiationirr:LineMA3 | 0.1554541 | 0.0361867 | 0.2749441 | 1000.0 | 0.010 |
| Irradiationirr:Day2 | -0.1234450 | -0.2509863 | -0.0002705 | 1000.0 | 0.052 |

[a]The intercept represents the control treatment with no other males but a female mating partner in the socio-sexual environment (group “mated”) in males from line S1.

<u>S5: Fertility reduction after short-term recovery period, statistics summary</u>

#### Final reduced model

Iterations = 200001:5195001
Thinning interval = 5000
Sample size = 1000
DIC: 2700.433

R-structure: ~idh(Irradiation:OtherMales:Mating):units

| | post.mean | l-95% CI | u-95% CI | eff.samp |
|---|---|---|---|---|
| Irradiationctrl:OtherMalesNoMales:Matingmated.units | 0.0005513 | 1.398e-07 | 0.00304 | 906.9 |
| Irradiationirr:OtherMalesNoMales:Matingmated.units | 0.0613105 | 4.113e-03 | 0.12081 | 673.2 |
| Irradiationctrl:OtherMalesOtherMales:Matingmated.units | 0.0654038 | 2.962e-02 | 0.10789 | 1000.0 |
| Irradiationirr:OtherMalesOtherMales:Matingmated.units | 0.0274842 | 1.702e-07 | 0.07281 | 700.9 |
| Irradiationctrl:OtherMalesNoMales:Matingvirgin.units | 0.0052644 | 1.021e-07 | 0.01544 | 859.2 |
| Irradiationirr:OtherMalesNoMales:Matingvirgin.units | 0.0741622 | 3.533e-02 | 0.12321 | 1000.0 |
| Irradiationctrl:OtherMalesOtherMales:Matingvirgin.units | 0.1169395 | 5.377e-02 | 0.18930 | 1000.0 |
| Irradiationirr:OtherMalesOtherMales:Matingvirgin.units | 0.0616642 | 1.706e-02 | 0.11275 | 1000.0 |

Location effects: Offspring ~ Irradiation * Mating + Irradiation * OtherMales + Irradiation * Line + Irradiation * Day

| | post.mean | l-95% CI | u-95% CI | eff.samp | pMCMC |
|---|---|---|---|---|---|
| (Intercept) [a] | 4.3412639 | 4.2904147 | 4.3865824 | 815.8 | <0.001 |
| Irradiationirr | -1.7189030 | -1.8381161 | -1.5970116 | 1000.0 | <0.001 |
| Matingvirgin | -0.0101969 | -0.0675079 | 0.0392527 | 1000.0 | 0.700 |
| OtherMalesOtherMales | -0.0733665 | -0.1406886 | -0.0001325 | 899.5 | 0.030 |
| LineMA3 | -0.0415929 | -0.0948666 | 0.0079605 | 895.6 | 0.116 |
| Day2 | -0.0197231 | -0.0643878 | 0.0358869 | 840.8 | 0.452 |
| **Irradiationirr:Matingvirgin** | **0.1212729** | **0.0105026** | **0.2207173** | **762.1** | **0.036** |
| **Irradiationirr:OtherMalesOtherMales** | **0.0709586** | **-0.0437706** | **0.1931525** | **1000.0** | **0.218** |
| Irradiationirr:LineMA3 | 0.3298935 | 0.2242613 | 0.4255065 | 568.1 | <0.001 |
| Irradiationirr:Day2 | -0.0948055 | -0.2053005 | 0.0078705 | 739.8 | 0.088 |

[a]The intercept represents the control treatment with no other males but a female mating partner in the socio-sexual environment (group “mated”) in males from line S1 on the first experimental day.

<u>S6: Fertility reduction after long-term recovery period, statistics summary</u>

#### Final reduced model

Iterations = 200001:5195001
Thinning interval = 5000
Sample size = 1000
DIC: 2720.99

R-structure: ~idh(Irradiation:OtherMales:Mating):units

| | post.mean | l-95% CI | u-95% CI | eff.samp |
|---|---|---|---|---|
| Irradiationctrl:OtherMalesNoMales:Matingmated.units | 0.0004953 | 1.692e-07 | 0.002413 | 824.0 |
| Irradiationirr:OtherMalesNoMales:Matingmated.units | 0.0878831 | 3.777e-02 | 0.147207 | 1000.0 |
| Irradiationctrl:OtherMalesOtherMales:Matingmated.units | 0.0005392 | 1.547e-07 | 0.002898 | 1114.4 |
| Irradiationirr:OtherMalesOtherMales:Matingmated.units | 0.0675683 | 2.403e-02 | 0.121629 | 1000.0 |
| Irradiationctrl:OtherMalesNoMales:Matingvirgin.units | 0.0013676 | 2.162e-07 | 0.006248 | 1000.0 |
| Irradiationirr:OtherMalesNoMales:Matingvirgin.units | 0.0148451 | 1.081e-07 | 0.043081 | 706.8 |
| Irradiationctrl:OtherMalesOtherMales:Matingvirgin.units | 0.0240508 | 9.667e-03 | 0.039511 | 844.3 |
| Irradiationirr:OtherMalesOtherMales:Matingvirgin.units | 0.0193200 | 2.323e-07 | 0.050879 | 452.9 |

Location effects: Offspring ~ Irradiation * Mating + Irradiation * OtherMales + Irradiation * Line

| | post.mean | l-95% CI | u-95% CI | eff.samp | pMCMC |
|---|---|---|---|---|---|
| (Intercept) [a] | 4.420200 | 4.382015 | 4.457470 | 885.9 | <0.001 |
| Irradiationirr | -1.571112 | -1.667179 | -1.482697 | 708.2 | <0.001 |
| Matingvirgin | -0.015869 | -0.055491 | 0.026464 | 911.6 | 0.460 |
| OtherMalesOtherMales | -0.025176 | -0.064166 | 0.012497 | 887.0 | 0.202 |
| LineMA3 | -0.018699 | -0.058589 | 0.020819 | 862.9 | 0.372 |
| **Irradiationirr:Matingvirgin** | **0.067459** | **-0.036501** | **0.148115** | **765.6** | **0.160** |
| **Irradiationirr:OtherMalesOtherMales** | **0.005423** | **-0.078215** | **0.090581** | **608.2** | **0.904** |
| Irradiationirr:LineMA3 | 0.164696 | 0.072948 | 0.252892 | 261.8 | <0.001 |

[a]The intercept represents the control treatment with no other males but a female mating partner in the socio-sexual environment (group "mated") in males from line S1.
